# Functional Mapping and Engineering of the Sec Translocon Unlocked by a Cell-Free System

**DOI:** 10.64898/2025.12.09.688994

**Authors:** Markus Meier, Scott A. Scholz, Leo von Bank, João E. Levandoski, Mosche Lückhof, Maximilian Schaaf, Shutian Si, Ingo Lieberwirth, Anna Lena Jung, Katharina Landfester, Arnold J.M. Driessen, Tobias J. Erb

**Affiliations:** Department of Biochemistry & Synthetic Metabolism, Max Planck Institute for Terrestrial Microbiology; Karl-von-Frisch-Str. 10, D-35043 Marburg, Germany; Institute for Lung Research, Universities of Giessen and Marburg Lung Center, Philipps-University Marburg, German Center for Lung Research (DZL); Hans-Meerwein-Straße 2, D-35043 Marburg, Germany; Core Facility Flow Cytometry - Bacterial Vesicles, Philipps-University Marburg; Germany; Department Physical Chemistry of Polymers, Max Planck Institute for Polymer Research; Ackermannweg 10, D-55128 Mainz, Germany; Department of Molecular Microbiology, Groningen Biomolecular Sciences and Biotechnology Institute, University of Groningen; Nijenborgh 7, 9747AG Groningen, Netherlands; Center for Synthetic Microbiology (SYNMIKRO), Philipps-University Marburg; Germany

## Abstract

Almost all proteins are inserted or translocated across membranes by the universally conserved Sec translocon. Despite its central role, experimental access to Sec function has remained limited. Here, we present a cell-free protein synthesis platform that inserts SecYEG into synthetic vesicles, enabling direct testing of Sec in real-time and high-throughput, circumventing longstanding viability constraints. Screening 300 Sec variants in a single experiment, we consolidate three decades of Sec research, while vastly expanding mutant diversity for structure-function insights. Mapping over 30 functionally critical regions that modulate Sec activity across three orders of magnitude, we uncover dozens of super-active variants. We further leverage our system to increase membrane protein quality and nanobody export, highlighting the potential of our system for advancing applications in synthetic biology and biotechnology.

## Introduction

Membrane proteins functionalize the phospholipid bilayer of cells and serve in various essential biological processes, including energy conversion, sensing, transport, light harvesting, motility and membrane stability (*1*). Intrinsically, membrane proteins are largely hydrophobic and tend to aggregate in the cytoplasm. Therefore, cells have evolved molecular machineries to facilitate their insertion into and translocation across the bilayer. The prime example is the Sec translocon (SecYEG in bacteria, SecYEβ in archaea, Sec61 in eukaryotes) (*2*). This heterotrimeric membrane complex consists of one of each subunit SecY, SecE and SecG and is essential to life (*3*). Homologs of the central pore protein SecY are found across all domains of life, and SecY can be even traced back to the last universal common ancestor of all cells (*4*).

SecYEG inserts transmembrane proteins into the bilayer in a co-translational fashion. The translating ribosome is directed to the translocon, which integrates the growing nascent chain into the membrane (*3, 5*). Beyond insertion, SecYEG also serves as primary protein export system. In bacteria, protein export takes place post-translationally and is powered by the cytoplasmic ATPase SecA (*6*). Hydrophobic alpha-helices in the form of transmembrane domains or N-terminal signal peptides act as recognition signals for SecYEG.

Despite decades of research, efforts to both understand SecYEG and engineer the pore for improved function for synthetic biology (e.g., construction of synthetic cells) or biotechnology (e.g., enhanced protein secretion) remains constrained. This is because the essentiality of SecYEG and its propensity for toxic ion-leaking mutants make it difficult to manipulate the pore *in vivo* or even produce it in living cells for *in vitro* assays (*3, 7, 8*). Despite the development of different methods, *in vitro* assays on SecYEG nonetheless still required complicated workflows that restrict both high-throughput and quantitative analysis (*9*).

*In vitro* cell-free protein synthesis (CFPS) offers a solution by removing viability constraints, which has been exploited to study many essential proteins (*10*). Critically, CFPS can directly insert membrane proteins into phospholipid bilayers (*11*), which has been leveraged to demonstrate insertion of functional SecYEG into empty membranes (*12, 13*). However, characterization of SecYEG translocation activity still remained semi-quantitative, relying on labor- and time-intensive methods based on protein substrate protection from proteolytic digestion, SDS-PAGE and fluorescence or [^35^S]methionine detection (*9, 12*).

To overcome these experimental limitations in studying and engineering the Sec pore, we established a CFPS-based platform that allows functionalization of blank membrane vesicles with SecYEG. Our platform enables evaluation of translocation and insertion activities of several reporters via quantitative high-throughput and real-time kinetic readouts (*14*), revealing insights about Sec and providing a convenient gateway for studying membrane and exported proteins. By directing protein substrates to the membrane, our system reduces aggregation during CFPS, increases translocation and promotes correct insertion orientation, which opens new avenues for studying and engineering of the basic functions of membrane proteins and construction of functionalized membranes from the bottom-up.

## Results and Discussion

### Establishment of an assay to quantify expressed SecYEG translocation activity

To produce SecYEG and quantify its activity *in vitro*, we paired CFPS of SecYEG in the presence of blank vesicles with an adapted split nanoluciferase-based translocation assay (*14*). In this assay, the SecYEG pore is produced *in vitro* from a plasmid (pSecYEG; Fig. S2a) and (self-)inserted into the blank membrane vesicles containing a catalytically inactive nanoluciferase fragment (11S). Translocation activity of SecYEG is probed by a plasmid-encoded translocation reporter protein that carries a C-terminal tag for nanoluciferase complementation (*15*). Upon expression and translocation of the reporter protein across the vesicle membrane, the C-terminal tag complements the 11S, which results in a luminescence signal upon enzymatic conversion of furimazine to furimamide (Fig. 1a-c).

**Figure 1.**
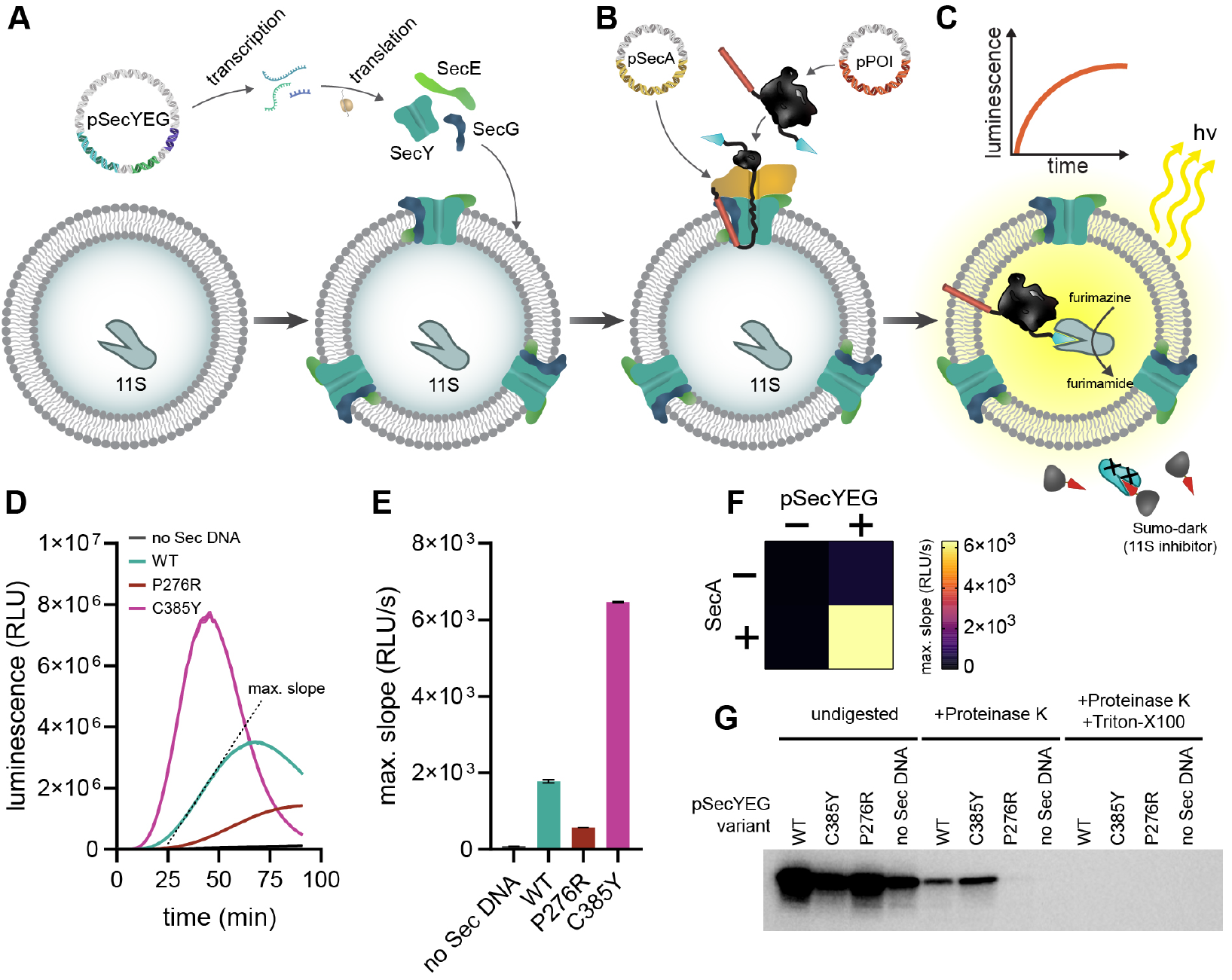
A SecYEG plasmid for cell-free expression establishes functional translocons on blank membranes. A) A plasmid encoding *E. coli secYEG* (pSecYEG) is expressed in CFPS leading to spontaneous assembly of functional translocons into blank reporter vescles. B) Powered by the co-expressed SecA ATPase, SecYEG complexes facilitate the translocation of newly synthesized reporter proteins into the vesicle lumen. C) Schematic representation of the real-time translocation assay based on split nanoluciferase. Only successfully translocated reporter proteins with a C-terminal complementation tag activate the encapsulated 11S nanoluciferase fragment leading to a luminescence signal. SUMO-dark proteins block 11S activity outside of the vesicles. D) Kinetic translocation data of proOmpA-pep99 reporter in PURE expression system using different SecY point mutants. Three encoded SecY variants were tested: WT, P276R and C385Y. E) The respective maximum (max.) slope values were used to quantify translocation activity. F) Heat map of pSecYEG-dependent translocation activity in the absence and presence of SecA (expressed separately and added during reporter expression). G) Luminescence blot of proOmpA-pep86 protected from Proteinase K digestion after SecYEG-dependent translocation into the vesicle lumen.

As translocation reporter substrate, we employed the first 178 amino acids of the precursor of the *E. coli* outer membrane protein (proOmpA), which is widely used as model for translocation (*16*). We fused proOmpA with a high affinity nanoluciferase complementation tag (pep86) at the C-terminus (proOmpA-pep86) and provided the gene on a separate plasmid. For vesicle encapsulation of the 11S nanoluciferase fragment, we chose a molar phospholipid ratio of 40% DOPC, 30% DOPE and 30% DOPG (*17, 18*). Co-expression of pSecYEG and pSecA, encoding the motor protein SecA, together with translocation reporter proOmpA-pep86 in a cell-free *E. coli* lysate resulted in a luminescent signal. In contrast, luminescence was strongly reduced for SecY mutants P276R, G240D and Y429D (*19-21*) that are known to be impaired in SecA-based translocation (Fig. S3a), establishing basic functionality and sensitivity of our assay.

To further improve the assay, we replaced the high affinity C-terminal tag with one of two lower affinity tags (pep99 and pep104), which we further call medium and low affinity tag, respectively (*15*). This reduced the overall signal, increasing the assay life time while also potentially opening up the upper dynamic range towards engineering highly active Sec variants (Fig. S1a,b; Fig. S2e,g). During reporter expression, we followed luminescence for 2 hours and used the maximum luminescence slope to quantify translocation activity. Compared to SecY WT, the P276R variant of SecY showed ∼5-fold reduction in translocation, while the previously described SecY C385Y mutant with improved translocation activity (*12*) showed a ∼7-fold increase in translocation activity in our *E. coli*-based cell-free *in vitro* system (Extended Data Fig.3c,d).

*E. coli* cell-free extracts still contain all proteins, including Sec components and auxiliary proteins. Thus, we switched to the PURE expression system, a CFPS comprised solely of purified components (*22*), to determine whether the expressed SecYEG and SecA alone were sufficient for translocation. The PURE-based assay mainly recapitulated the results of the *E. coli* system (Fig. 1d-f), and was independently confirmed through a Proteinase K protection assay (Fig. 1g). Compared to the SecY WT, P276R and C385Y showed ∼5-fold reduced translocation efficiencies, while the C385Y variant performed ∼4-times better (Fig. 1e). However, we noticed that the absolute luminescence signal in the PURE system was an order of magnitude higher than in S50 autolysate, which we traced down to the presence of native membrane vesicles in the autolysate that did not contain the 11S protein, thus lowering the luminescence signal (Fig. S4). We used this insight to further tune the dynamic range of our translocation assay in PURE by adding 3 mg/mL blank liposomes without 11S as “competitor vesicles” for all subsequent assays (Fig. S2b,c), establishing a robust, quantitative readout for SecYEG translocation.

### Development of an assay to quantify SecYEG-dependent membrane protein insertion

Having established and optimized an *in vitro* assay to track SecYEG translocation in real-time, we sought to develop a reporter to probe insertion of integral membrane proteins, which has been challenging, thus far. We selected the seven transmembrane domain protein proteorhodopsin (PR) that is inserted through SecYEG (*23*). We deleted the last transmembrane domain of PR and fused the low affinity complementation tag to its C-terminus (PR(-1tm)-pep104), so that the tag would become exposed to the vesicle lumen, complementing the 11S fragment, resulting in a luminescent signal (Fig. 2a).

**Figure 2.**
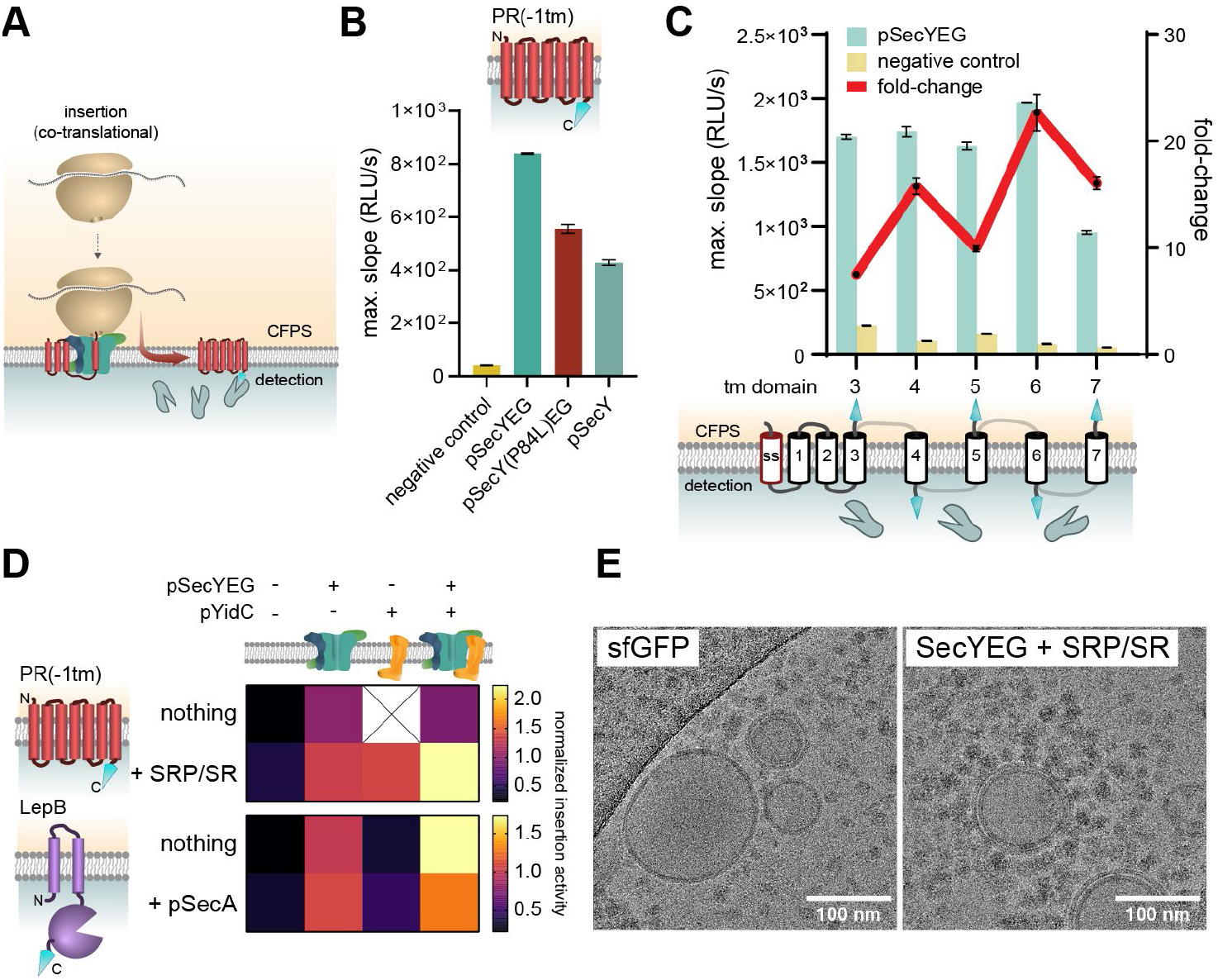
Membrane proteins produced in CFPS are directionally incorporated via pre-synthesized SecYEG. A) Schematic representation of the co-translational insertion of integral membrane protein reporters via SecYEG. The model membrane protein proteorhodopsin was used as a reporter (PR(-1tm)-pep104). The C-terminal transmembrane (tm) domain was removed so that the complementation tag faces the detection side of the reporter vesicles. B) Maximum slope data for insertion assays via pre-expressed SecYEG. C) Assessment of directional insertion of membrane proteins via SecYEG. Different transmembrane helices were removed iteratively from the C-terminus of proteorhodopsin and the resulting insertion signals measured in the luminescence assay. The fold-change of the signal with SecYEG was plotted against the signal for the transmembrane protein negative control. Correctly oriented membrane proteins should produce an alternatingly high/low insertion signal reflecting the C-terminus alternating between the detection and the CFPS side. D) Influence of auxiliary subunits on insertion of PR(-1tm)-pep104 and LepB-pep86. pYidC and pSecA were co-expressed with pSecYEG. The protein components or SRP (Ffh) and SR (FtsY) were pre-expressed in separate PURE reactions and added to the CPFS reaction expressing pSecYEG. Resource competition was compensated with expression of sfGFP as negative controls. E) Cryo-TEM micrographs of reporter vesicles after sfGFP (left) or SecYEG expression (right). Vesicle preparation by extrusion yielded predominantly uni- and bilamellar vesicles around 100 nm.

Integration of the PR reporter was dependent on SecYEG, with a 20-fold increased luminescent signal over the negative control (expression of a multi-pass transmembrane protein instead of SecYEG) (Fig. 2b). Compared to SecYEG WT, SecY variant P84L, which is impaired in insertion (*24*), or SecY alone (without SecE and SecG) showed about 60% and 50% luminescence, respectively, supporting the central role of SecY in insertion. When no SecYEG or multi-pass transmembrane protein was pre-synthesized, the PR reporter showed very high production levels with some spontaneous membrane insertion (Fig. S5a,b), which is in line with earlier findings (*11*). However, under these conditions, the unassisted insertion of PR resulted in significant aggregate formation and vesicle clumping. This finding was independently confirmed by dynamic light scattering (DLS) and multi-angle-DLS measurements, which demonstrated that SecYEG drastically improves protein expression quality (Fig. S6).

We also examined insertion directionality (*25*) by sequentially deleting C-terminal transmembrane domains of the PR reporter (Fig. 2c). This resulted in a strongly alternating luminescence pattern, when normalized against the negative control. This pattern corresponded to alternation of the C-terminus between the expression and the detection side of the membrane, indicating correctly folded and correctly inserted protein. Together, this data showed that PR is spontaneously integrated to a very basal level, but that SecYEG drastically improves insertion (∼20-fold), directionality (∼2-fold) and reduces aggregation.

### Auxiliary factors improve insertion

Next, we tested the impact of auxiliary factors, which are known to enhance and/or aid in SecYEG-dependent insertion, especially that of more complex substrates (*3*). We chose three known auxiliary factors of SecYEG, which we co-expressed or provided as pre-expressed protein to our *in vitro* system: SecA, YidC, and the signal recognition particle (SRP) / SRP receptor (SR) pathway. YidC is a membrane protein insertase that inserts integral membrane proteins individually or in concert with SecYEG (*26*). The SRP/SR pathway facilitates sorting of co-translationally integrated proteins to both SecYEG and YidC and consists of the two protein components Ffh and FtsY, as well as the 4.5S RNA (*5*). Because the 4.5S RNA is already part of the CFPS tRNA mix (*27, 28*), we did not provide or express additional RNA.

When SRP/SR was included in the reaction mixture, PR(-1tm)-pep104 integration was improved by 30% (Fig. 2d). Moreover, SecYEG and YidC showed a synergistic behavior, but only in the presence of SRP/SR, resulting in a more than 2-fold increased integration signal over SecYEG alone. This observation supports the finding that SecYEG and YidC form an insertase complex (*29*), which resembles the eukaryotic ‘multipass translocon’ and is independent of the lateral gate (*30, 31*). We also tested these factors alone and in combination with an alternative reporter protein, the signal peptidase LepB (LepB-pep86), which has been proposed to require SecYEG, SecA, as well as YidC for efficient insertion through temperature-sensitive mutants and cross-linking experiments (*32, 33*). We observed ∼70% increased LepB insertion when YidC was co-expressed (Fig. 2d), while the motor protein SecA alone, or in combination with YidC, did not further increase the signal, indicating that SecA was not essential for LepB insertion. In summary, these data showed that our *in vitro* assay is capable of assessing the contribution of different auxiliary factors to the insertion of specific membrane proteins that would be impossible with state-of-the art *in vivo* experiments.

### 294 SecY mutant library survey to quantify SecYEG activity

Our *in vitro* assay opened up the possibility to assess the translocation and insertion activities of SecYEG variants in minimal volume, in parallel, and at high-throughput. Therefore, we set out to validate our method against 70 published mutants, while characterizing 204 new variants, and 20 mutants with *secE* and *secG* deleted in a single experiment (Data S1). To array, quantify and distribute DNA templates, we established a contact-free liquid handling workflow (Fig. S7a). Translocation data for the SecY library was collected in triplicates on separate days and showed a reproducible, robust output (Pearson’s r between replicates 0.889 – 0.957) (Fig. 3b).

**Figure 3.**
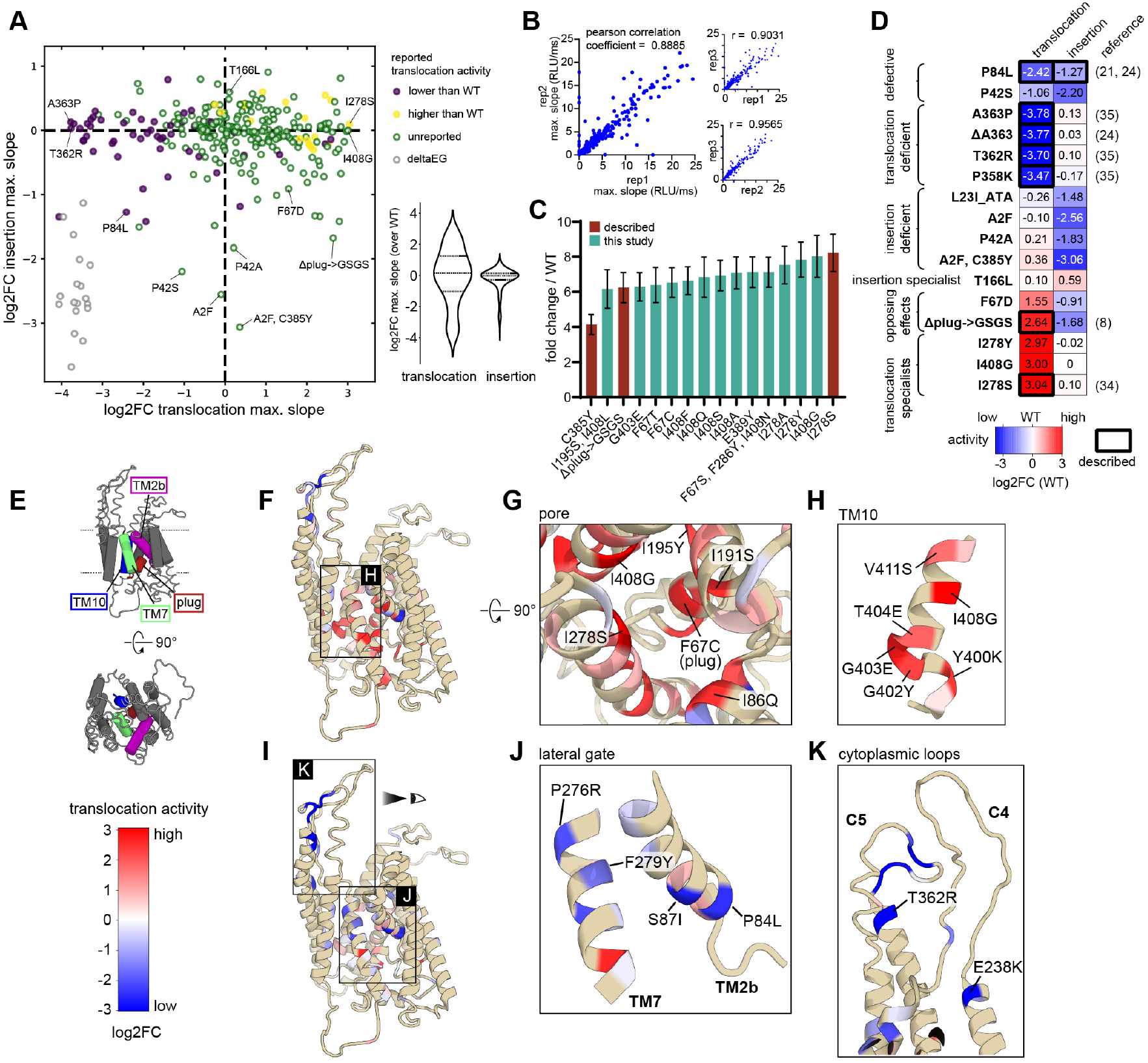
Measurement of translocation and insertion activities of a 294-variants SecY library. A) Insertion activity plotted against the translocation activity of each variant as log2-fold change compared to SecY WT (dashed lines). Previously described SecY variants with the translocation activity reported higher than WT are shown as yellow, variants with lower reported activity are shown as purple dots. New variants are shown as green and deltaEG variants (*secE* and *secG* deleted from pSecYEG) as grey circles. B) Correlation between replicates of the three separate translocation experiments. C) Top 5% translocating mutants of the SecY library shown at fold-change over WT activity. D) Heatmap of selected SecY library variants with special combinations of translocation and insertion activity. Bold boxes mark variants which have been described previously. E) Schematic representation of the SecY structure. Highlighted are TM2b (pink), plug (*8*) (red), TM7 (green) and TM10 (blue). F) Point mutants with the highest translocation activity obtained for each probed residue are plotted on the SecY structure according to their log2-fold-change over the WT activity. G) Cytoplasmic view into the SecY central pore highlighting the plug and the pore ring residues I86, I191, I278 and I408. H) Zoom into TM10 which displays a high propensity for super-active mutants. I) Point mutants with the lowest translocation activity obtained for each probed residue. J) Zoom into the lateral gate helices TM7 and TM2b. K) Zoom into the cytoplasmic loops C4 and C5, responsible for ribosome and SecA binding. All values were plotted on a previously published structure (*23*) (PDB: 5ABB).

Note that the activity of SecYEG variants described in the literature was mainly of qualitative nature, thus far. Moreover, previously described SecY variants could not be easily compared to each other, as they were examined in different experimental setups with various methods, substrates and even different signal sequences. Our data collected on the SecY library captured most of the literature findings (78% agreement) in respect to general trends, i.e., increased or decreased translocation activity (Fig. 3a; Data S1). Critically, our screen also provided detailed quantitative insights, revealing that the different SecYEG variants span a broad range of translocation activities from about 16-fold reduction to 8-fold increase (Fig. 3a).

Beyond confirming C385Y as highly-active translocation variant (*12*), we identified 77 additional mutants in 39 different sites with at least 2-fold and up to 8-fold improved activity over WT, including I278S (*34*) and novel variants I278Y, I278A, E389Y, G403E, I408G and I408A (Fig. 3c). Thereby, within one experiment, we were able to consolidate three decades of SecYEG research and examine previous and novel SecY variants on a directly comparable base (Fig. 3d;Data S1). This approach enables an unprecedentedly improved throughput (∼2 orders of magnitude (*21, 24, 35, 36*)) by directly testing hundreds of genetic constructs side-stepping transformations, protein purification and *in vitro* reconstitution, while providing high resolution and large dynamic range.

In SecY, different structural domains are connected to distinct functions like targeting (*3, 5*), proofreading (*37*) and SecA/ribosome binding (*38-40*). To elucidate this structure-function relationship, we mapped our data on a SecY structure (PDB: 5ABB). Mutants in the C4 and C5 loops led to severe translocation defects (Fig. 3k), which is in line with the role of these sites as major contact points for the ribosome and SecA (*38-40*). Conversely, several known (F67 (*41*), I195 (*36*), I278 (*34*), G403 (*24*) and I408 (*36, 42*)) and new positions (F64, L72, M83, I86, L285 and G402) within the transmembrane regions of SecY gave rise to super-active mutants, with TM10 being a major hotspot (Fig. 3h; Fig. S7). Especially striking was the high frequency of super-active translocators generated by diverse mutations of the hydrophobic pore ring of SecY, which conveys proof-reading of translocated substrates via conserved isoleucines (*37*) (Fig. 3g).

The full SecYEG library was also evaluated for aiding integration of a model membrane protein. In contrast to the broad range of observed translocation activities (spanning 138-fold, standard deviation 3.5-fold), we observed a smaller impact of mutations on insertion activity (spanning 36-fold, standard deviation 1.7-fold) (Fig. 3a), most of them with decreased activity. Interestingly, several of these variants were not impacted in translocation, showing that the translocation and insertion pathways are under (partially) different constraints (Fig. 3d; Fig. S8c,d).

However, we identified one novel insertion variant, T166L that showed ∼30% increased insertion activity but WT-like translocation (Fig. 3d), which we validated in an independent experiment (Fig. S8b,d). Interestingly, T166 is a rare polar amino acid within the transmembrane region of TM4 and lies just beneath the hydrophilic TM3-4 groove, a region which has recently been characterized to have chaperone activity during insertion (*43*). In summary, our data provide for the first time a comprehensive and comparable overview about the translocation and insertion activities of more than 300 SecYEG variants, providing quantitative, high-quality data for 70 published mutants and identifying new variants with completely novel properties.

### Identifying signal peptides for efficient translocation of a nanobody

Our library showed that mutations of the central SecY pore globally alter translocation activity of SecYEG itself. However, at the level of individual substrates, signal peptides influence how readily and through which exact pathway a given substrate is translocated (*44, 45*). We surmised that our *in vitro* system could be leveraged to screen for secretion of biotechologically-relevant proteins. To test this possibility, we screened a set of 11 signal peptides for translocation of an anti-mouse-IgG nanobody (TP1170) (*46*), as an orthogonal Sec substrate. We chose native signal peptides from *E. coli* (*44*) and *C. glutamicum* (*47*), as well as synthetically derived sequences (*47-49*) (Fig. 4c; Tab. S1), which we N-terminally fused to the nanobody carrying a C-terminal high affinity complementation tag (Fig. 4b). Notably, translocation activities ranged almost 70-fold between the weakest (PorB) and the strongest signal sequence, OmpA(extCore) (Fig. 4d). The translocation of the strongest two peptides, OmpA(extCore) and YncJ, was even further improved ∼3-fold when swapping SecY WT to the super-active SecY(I408G) variant that we had identified earlier in our library screening (Fig. 4e). Altogether, these experiments demonstrate that our *in vitro* system is able to identify signal peptide-substrate combinations that can work synergistically with improved SecYEG pores, allowing to tackle both the Sec machinery and substrate proteins for improved translocation.

**Figure 4.**
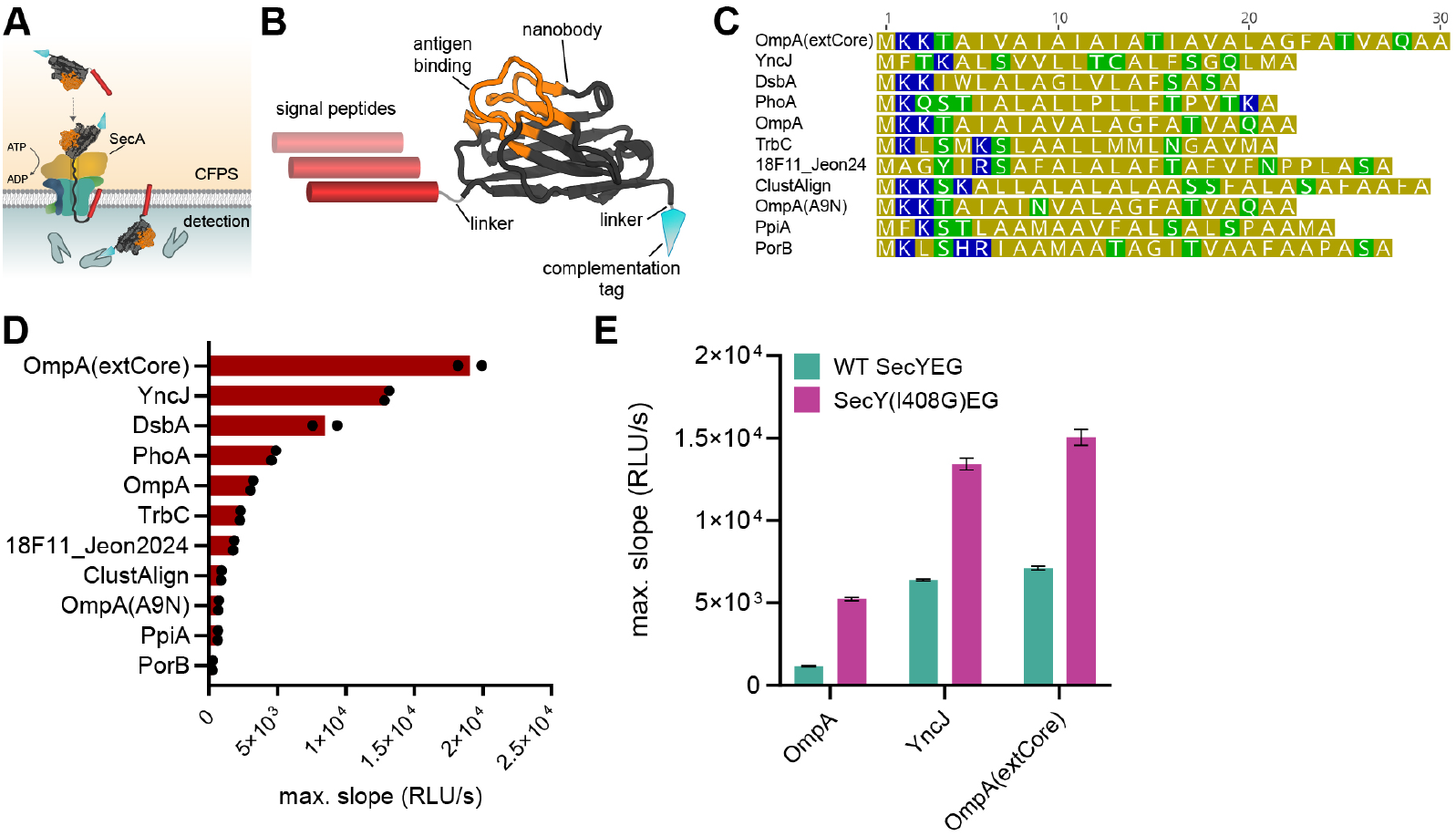
Screening signal peptides to translocate a nanobody through SecYEG. A) Schematic of the *in vitro* setup to measure SecYEG- and SecA-dependent nanobody translocation using the luminescence assay. B) Structural representation of the anti-mouse-IgG nanobody (TP1170) reporter with an N-terminal signal peptide and a C-terminal complementation tag. The nanobody structure was predicted using AlphaFold Server. All reporters were encoded in separate plasmids. C) Overview of the amino acid sequences of the 11 tested N-terminal signal peptides to translocate the nanobody through SecYEG. D) Translocation activities of the nanobody constructs with different signal peptides. SecYEG and SecA were both expressed from plasmids to promote post-translational translocation. E) Translocation assays of the nanobody combining the best two signal peptides, YncJ and OmpA(extCore), and the super-active pSecY(I408G)EG variant identified in the library screening. For comparison, the nanobody with the OmpA signal peptide was used.

## Discussion

SecYEG is the central hub for protein partitioning and strictly essential for living cells, which makes it a keystone research subject, but also hard (to impossible) to assess the function of SecYEG mutants *in vivo*. In this study, we adapted and implemented a cell-free system to synthesize, study and engineer functional SecYEG and its associated components. This cell-free approach decouples activity from cell survival, which allows to characterize even potentially lethal (i.e., non-active or super-active) variants in real-time and at high-throughput. In the future, screening of SecYEG will be only limited by DNA synthesis and liquid handling. Furthermore, the bottom-up reconstruction of the SecYEG pore in blank vesicles allows to conveniently assess the effects of auxiliary Sec components, such as YidC, SecA and SRP/SR, which opens up the possibility to study the complex interplay of these proteins with the pore without the risk of pleiotropic effects.

Gathering translocation and insertion data of an extensive library of SecY mutants, we expanded the list of SecY variants described over last three decades by almost fourfold, and identified several new variants with increased translation, but notably, for the first time, also insertion activity. In our screen of ∼300 variants, we observed 77 variants with at least 2-fold increased translocation activity covering almost 40 different sites in SecY. Even though we targeted specific sites of the protein, this surprisingly high likelihood to produce a (super-)active translocator indicates that SecY did not evolve for maximum translocation activity, but rather membrane tightness and proof-reading (*50*). The relatively rare occurrence of improved insertion variants raises the question of whether insertion activity has been evolutionarily optimized or rather under sampled by the field, due to a bias towards translocation activity.

Our work lays the proof of principle for the directional insertion and production high quality membrane proteins and for the efficient translocation of biotechnologically-relevant proteins. These insights will be essential for future efforts to improve membrane protein production in natural and artificial cell systems. Together, our approach alleviates longstanding challenges in Sec research, laying a foundation to better understand the Sec machinery, and enables synthetic biology and biotechnology applications (*51*).

## Supporting information

Supplementary Data S1 - pSecYEG library

Supplementary Data S1 - Plasmids

## Acknowledgments

We would like to thank Dr. Tom Robinson and Dr. Naresh Yandrapalli for early discussions and technical support. We thank Dr. Amir Pandi for instructions on autolysate production, and Dr. Blake Rasor for for pJL1-sfGFP plasmid and instructions on lysate production. We thank Dr. Sebastian Barthel for the sfGFP-pepper plasmid and help with transcription/translation measurements. We thank Dr. Frank Abendroth for the pET29-LgBiT plasmid, and we gratefully acknowledge the support and resources provided by the ChemBio research-based platform at Marburg University. We thank Dr. Andreas Küffner for providing us with the SUMO plasmid. We thank Dr. Owen Jarman, Elisabeth Bobkova, Dr. Nataliya Safronova, Dr. Blake Rasor, Dr. Beau Dronsella, Max Hoffmann-Becking, Dr. Max Gantz, Nitin Bohra and Dr. Martine Ballinger for regular discussions, ideas and feedback on the manuscript.

## Funding

Gordon and Betty Moore Foundation, GBMF10652 (grant DOI https://doi.org/10.37807/GBMF10652)

Merck Future Insight Prize

AJMD is supported by the Netherlands Organization for the Advancement of Science (NWO) through the ‘BaSyC – Building a synthetic cell’ Gravitation grant (024.003.019) of the Ministry of Education, Culture, and Science.

## Author contributions

All authors contributed to the design, execution, interpretation of the experiments, and writing and revision of the manuscript.

SAS and TJE conceived the work.

MM, SAS and TJE wrote the manuscript.

MM led the experimental work, including the development and implementation of the luminescence assay system, translocation and insertion assays, data analysis, figure compilation, experimental design, and library design.

LvB contributed to insertion assays with auxiliary components, and designed and performed nanobody translocation assays.

JEL investigated the influence of native vesicles in lysate on translocation assay performance.

SAS performed protease protection assays, and participated in library design and constructed the pSecYEG plasmid.

ALJ analyzed nano-FCM measurements.

ML performed and analyzed the nano-FCM measurements. KL analyzed DLS measurements of vesicles.

MS performed and analyzed DLS measurements of vesicles.

SS conducted cryoTEM imaging and analysis of cryoTEM data, with IL providing additional data analysis.

AJMD advised on strategic and experimental design. Funding acquisition: ALJ, AJMD, KL, SAS, TJE

All authors contributed to the writing and final approval of the manuscript.

## Competing interests

The authors declare no competing interests.

## Data and materials availability

All data are available in the main text or the supplementary materials. Key plasmids are available on Addgene.

## Notes

### Competing Interest Statement

The authors have declared no competing interest.

