## Supplementary information for "Functional Mapping and Engineering of the Sec Translocon Unlocked by a Cell-Free System"

**The PDF file includes:**

Materials and Methods
Figs. S1 to S8
Table S1
References

**Other Supplementary Materials for this manuscript:**

Data S1 to S2

### Materials and Methods

Chemicals and reagents

The library was ordered from GenScript, primers were ordered from Eurofins and gBlocks from TWIST Biosciences. Lipids were ordered as chloroform solutions (Avanti). The Nano-Glo HiBiT Extracellular Detection System, The Nano-Glo HiBiT Blotting System and QuantiFluor dsDNA System were purchased from Promega. For plasmid cloning, Q5 High-Fidelity DNA Polymerase 2x Master Mix, NEBuilder HiFi DNA Assembly Master Mix, DpnI, KLD Enzyme Mix and NEB Turbo *E. coli* were supplied by New England Biolabs. The NucleoSpin Plasmid miniprep and PCR clean up kits were supplied by Macherey-Nagel. PURE CFPS systems were purchased from Hölzel Diagnostika Handels GmbH (PUREfrex 1.0, Gene Frontier) or from New England Biolabs (PURExpress). Murine RNase inhibitors were purchased from New England Biolabs. Amino acids and tRNAs for lysate reactions were supplied by biotechrabbit (RTS Amino Acid Sampler) and Roche Diagnostics (tRNA, from E.coli MRE 600), respectively. Enzymes and reagents which were indicated for storage in ultra-low temperature freezers (including PUREfrex and PURExpress) were stored at -70°C to reduce energy consumption.

Construction of pSecYEG

To synthesize the three *E. coli* Sec core proteins, SecY, SecE and SecG, we constructed a genetic base construct, the pSecYEG plasmid. It was designed for high expression of the core *E. coli* translocon from a single operon (Fig. S2a). To ensure compatibility with common *E. coli* CFPSs and avoid toxicity issues during cloning we used the T7 promoter. While the relative expression levels were not optimized for translocation activity, we tuned the translation initiation rate of the core components to match the SecYEG translocon stoichiometry (*52*). The operon was concluded with two consecutive terminators: L3S2P21 (*53*) and T7 terminator.

Protein purifications of SUMO-dark and 11S

SUMO-dark and 11S proteins were purified via N/C-terminal His-tags. Plasmids under T7 control were transformed into chemically competent BL21 (DE3) cells. The whole plate of transformants was used to inoculate 1 L TB medium with the appropriate antibiotics and grown in 5 L baffled Erlenmeyer flasks at 25 °C for 24 h. The cells were not induced as the leakiness of the cells ensured high levels of protein production. Cells were harvested by centrifugation and resuspended in buffer D (20 mM Tris-HCl pH7.5, 50 mM KCl) + 20 mM imidazole supplemented with 10 µg/mL DNase I (Thermo Fisher) and 0.1 mM PMSF. Cells were lysed using a homogenizer (EmulsiFlex-B15, Avestin) and proteins extracted from the cleared lysate using Protino Ni-NTA agarose in gravity flow columns (Protino, Machery-Nagel). Proteins were eluted in buffer D + 500 mM imidazole and the imidazole removed using PD-10 desalting columns (Sigma-Aldrich) eluting into buffer D. Proteins were concentrated using amicon spin concentrators (3 kDa MWCO, Sigma-Aldrich), snap-frozen in liquid nitrogen and stored at -70 °C.

Liposome preparation

Large unilamellar vesicles (LUVs) were prepared by lipid swelling and subsequent extrusion. 5 mg phospholipids in chloroform were added to a glass test tube and dried under a stream of nitrogen followed by applying vacuum for at least 1.5 h. The standard lipid composition was DOPC:DOPE:DOPG 40:30:30 molar ratio. The vacuum was released with nitrogen and 500 µL of inner aqueous solution (IA) added at room temperature. 500 µL of IA contained 51 µL 11S protein (LgBiT, Nano-Glo HiBiT Extracellular Detection System, Promega) and 12µL 50% glycerol (w/v) in 17.5 mM HEPES pH7.2. Alternatively, to the commercial LgBiT stock, 7 µM purified 11S protein was used. In that case, 51 µL 50% glycerol (w/v) was added instead of the commercial LgBiT stock to adjust osmolarity. The suspension was incubated at room temperature for 30 min with occasional vortexing. After 5 freeze-thaw cycles between liquid nitrogen and a room temperature water bath, the suspension was extruded 11 times through a 0.1 µm polycarbonate membrane (GE Healthcare) using a mini-extruder (Avanti Research). Liposomes were washed 3 times by centrifugation for 1h at 45,000 g and 4 °C, removing the supernatant and resuspension in liposome wash buffer (6.3% glycerol, 17.5 mM HEPES pH7.2). Finally, reporter LUVs were resuspended in 50 µL liposome wash buffer yielding about 42 mg/mL lipids and stored at 4 °C for up to 5 days.

Empty "competitor" LUVs without encapsulated proteins were prepared similarly but instead, 5 mM HEPES pH7.2 was used as IA and for resuspension while freeze-thawing was omitted. Empty LUVs were concentrated to 60 mg/mL by centrifugation, snap-frozen in liquid nitrogen and stored at -70 °C.

Nanoluc translocation and insertion assays

For real-time kinetic translocation assays, a nanoluc based in vitro assay was adapted for the use in our system (*14*). Briefly, reporter liposomes contain the catalytically inactive 11S (LgBiT) variant of nanoluc which is activated by a high affinity complementation tag (pep86) or alternatives of it (*15*). Upon successful translocation of a reporter protein with the C-terminal complementation tag into the liposome lumen, a luminescent signal is emitted and detected in a plate-reader. To eliminate unspecific luminescence due to liposome breakage, a protein tagged with the dark peptide, in our case SUMO-dark, is added to the bulk solution (*14*).

To ensure proper translocon activity, the assay was divided into two phases: first the expression of the Sec system, followed by reporter expression and luminescence recording. The primary CFPS containing 60-150 µM SUMO-dark, 0.2 nM pSecYEG, 0.1 nM pSecA, 3.3 mg/mL blank competitor vesicles (without 11S; not added in lysates) and glycerol to adjust osmolarity (only lysates; measured with OM807, Loeser) was assembled on ice. Per 10 µl CFPS, 1 µL 42 mg/mL reporter liposomes were added on top and incubated in PCR tubes for 3h at 30 °C. From this point, samples were not cooled beneath room temperature to avoid precipitation of hydrophobic proteins. In the meantime, reporter CFPS was prepared: 60-150 µM SUMO-dark, 0.4 nM reporter DNA, glycerol to adjust osmolarity (only in lysates) and 1x furimazine (Promega). After 3h of Sec expression, primary and reporter CFPS were mixed at room temperature at equal ratio, and 9 µl each were dispensed into a 384-well plate (#784904, Greiner) in technical triplicates. The plate was sealed with transparent cover foil, centrifuged briefly (400 g) and immediately measured in a plate-reader (Infinite 200 Pro, Tecan) at 30 °C.

Insertion of integral membrane proteins was measured according to the same principle as translocation. In this case, SecA was omitted. To account for CFPS resource competition between the reporter and the Sec constructs, non-labelled full-length proteorhodopsin (pTE5620) was expressed instead of pSecYEG as a negative control plasmid that does not convey insertase activity. If applicable, membrane bound auxiliary components, like the YidC insertase, were co-expressed with pSecYEG at 0.1 nM plasmid concentration. Soluble components, like SRP and SR, were usually expressed separately and added at a ratio of 1:20.

SecY library translocation and insertion assays

Translocation and insertion activities for the library of pSecYEG variants were examined in luminescence assays. The library was ordered as arrayed clonal and purified plasmids (Genscript), dissolved in water and stored in a 384-well Echo source plate (384pp, Beckman Coulter). The concentration of each variant was measured with the QuantiFluor dsDNA System (Promega) by transferring 1 µL each into wells of 96-well plates using an Echo 525 liquid handler (Beckman Coulter), adding 200 µL QuantiFluor reagent and measuring fluorescence in a plate-reader (Infinite 200 pro, Tecan). Assay plates were prepared by transferring 0.6 fmol DNA of each variant into separate wells of a 384-well plate (#784904, Greiner) using an Echo 650T liquid handler (Beckman Coulter). A sufficient amount of assay plates for all experiments and replicates was prepared at the same time, sealed with adhesive PCR plate foils and stored at -20 °C.

For a luminescence assay, one plate containing all variants and control plasmids was thawed, the cover foil removed and the water evaporated by applying vacuum for 15 min using a desiccator. On ice, 3 µL primary CFPS containing all necessary components (except pSecYEG) including reporter liposomes was added to each well using a multi-channel dispenser (VOYAGER, Integra). For translocation experiments, the mix contained 0.1 nM pSecA. This resulted in a pSecYEG concentration of 0.2 nM in the primary CFPS. The plate was sealed with a transparent cover foil, mixed by orbital shaking, briefly centrifuged at 400 g and then incubated at 30 °C for 3 h. After this, 3 µL reporter CFPS was added per well. The plate was sealed, mixed, centrifuged and luminescence kinetics immediately measured for 2 h at 30 °C in a pre-heated plate-reader (Infinite 200 Pro, Tecan). Reporter CFPS contained 0.4 nM proOmpA-pep99 (translocation) or 0.4 nM PR(-1tm)-pep104 reporter DNA (insertion).

Library assay replicate data for translocation and insertion was all collected on separate days and processed. At each collected time point a luminescence slope was calculated with a linear regression including the previous and subsequent time point (3-point calculation). The maximum raw luminescence (trans/insert_highest_value) and slope value (trans/insert_peak_slope) are reported in (Data S1). In order to correct for assay variance between days, the average global peak_slope for every library member was divided by the average of the global peak_slope of all replicates (resulting in a minor adjustment since the daily variance was low). Rarely, there was the appearance of an outlier replicate which was removed from the normalized data, defined by a >25% difference from the average of all replicates, resulting in removal of <10% of normalized values. The normalized average was recalculated using the filtered values. This filtering step was not performed for the insertion data because there were 2 replicates. Finally Log2 fold-change was calculated compared to R22R_CGC for the WT reference. Please refer to (Data S1) for all raw and normalized values.

<u>Protease protection assays</u>

The successfully translocated and inserted reporter proteins were shielded form proteolytic digest and confirmed in protease protection assays (*9*). CFPS (15 µL) with 0.5 nM pSecYEG and 0.25 nM secA was assembled on ice and mixed with 1.5 µL 42 mg/mL freshly prepared and osmolarity matched empty liposomes (without 11S). The mix was incubated at 30 °C for 3 h and then, at room temperature, topped up with 15 µL fresh CFPS containing 0.2 nM reporter DNA (with pep86 tag). The reaction was incubated at 30 °C for 2.5 h to ensure successful translocation, transferred on ice and then gently mixed with 1x CutSmart (NEB) osmotically matched with glycerol (measured with OM807, Loeser). The mix was incubated on ice with 100 µg/mL Proteinase K (PK) and then precipitated with 7.5% ice cold TCA (w/v). As controls, samples were either not treated with PK or with PK and 1% Triton X-100. The precipitated samples were incubated on ice for another 30 min and then pelleted for 10 min at 17,000 g and 4 °C. After removing the supernatant, the pellet was washed with 500 µL ice cold acetone and centrifuged again and the supernatant was removed. The pellets were dried for 5 min at 37 °C, dissolved in 20 µL 2x Laemmli buffer and stored at -20 °C. After SDS PAGE, the proteins were transferred on a nitrocellulose membrane and visualized using the Nano-Glo HiBiT Blotting System (Promega) and (ChemiDoc MP, BioRad).

Dynamic light scattering (DLS) analysis of liposomes

The distribution of hydrodynamic diameters and polydispersity index (PDI) of liposomes after translocation and insertion reactions were analyzed using a Zetasizer Nano S90 (Malvern Panalytica GmbH). 10 µL sample were mixed with 190 µL PBS and measured in three technical replicates. Z-average values were calculated as averages and derivations of the main peaks.

Cryogenic transmission electron microscopy (cryo-TEM) of liposomes

Cryo-TEM micrographs of liposomes were acquired using a FEI Titan Krios G4 TEM (Thermo Scientific). 3 µL sample were applied to a Quantifoil or lacey carbon coated TEM grid which had been glow discharged in a Diner Nano oxygen plasma cleaner (Diener electronic) briefly before. Excess sample solution was removed with a filter paper and the grid plunged into liquid ethane using a Vitrobot Mark V (Thermo Fisher). Imaging was done under cryogenic conditions with an acceleration voltage of 300 kV. Micrographs were acquired under low dose conditions using a 4k Direct Electron Detection Camera (Gatan K3).

Lysate preparation

Cell extracts for lysate based cell free expression were prepared from BL21 Gold (DE3) cells either following a protocol for disruption by homogenization (*54*) or autolysis (*55*). In short, cells were grown to mid-log phase, harvested by centrifugation and resuspended in S30A buffer followed by lysis using a high-pressure homogenizer (EmulsiFlex-B15, Avestin) or by freeze-thawing and vortexing for cells bearing the pAD-LyseR plasmid. Cell debris was removed by centrifugation for 1h at 12/30/50/80 or 105k g. Autolysates were subjected to a run-off at 37°C for 2h before snap-freezing and storing lysates at -70 °C.

Cell free lysate reactions

Apart from using PURE kits, cell free reactions were performed in *E. coli* lysate based CFPS systems. The energy mix containing all necessary factors to ensure high expression was prepared as previously described (*56*) with some optimizations (*57*). Cell free reactions were assembled on ice with the final composition being: 33% autolysate or 27% homogenizer extract, 0.5 mM aa'S (except 0.42 mM Leu), 53 mM HEPES, 1.04 mM ATP and GTP, 0.63 mM CTP and UTP, 0.040 mg/mL tRNAs, 0.13 mM CoA, 0.17 mM NAD, 0.57 mM cAMP, 0.068 mM folinic acid, 1.00 mM spermidine, 25 mM 3-PGA, 0.67 U/µL murine RNase inhibitors (NEB). Mg-glutamate, K-glutamate, DTT and PEG were individually titrated for each lysate. Usual concentrations were: 6 mM Mg-glutamate, 80 mM K-glutamate, 1.5 mM DTT and 3% PEG-8000. DNA was added between 0.1 and 1 nM but usually 0.2 nM. For constructs based on expression from the T7 promoter, 0.25 µL T7 polymerase (NEB) was added per 10.5 µL reaction. Reactions were incubated in PCR tubes at 30 °C. For Fluorescence and luminescence measurements, the reactions were transferred into 384-well plates (#784076 or #784904, Greiner) and measured in a plate reader (Infinite 200 Pro, Tecan).

Measurement of transcriptional and translational activity of *E. coli* lysates

Transcription and translational outputs of *E. coli* cell free lysates were measured expressing a fusion construct encoding sfGFP and Pepper aptamer (*58*). Standard CFPS reactions with 1 nM pTE5422 (Addgene ID 229031) were assembled on ice and supplemented with 10 µM HBC620 (MedChemExpress, Monmouth Junction, NJ, USA, catalog no.: HY-133520). 10 µL reactions were transferred into 384-well plates (#784076, Greiner), sealed with a transparent cover foil, centrifuged briefly at 400 g and incubated in a pre-heated plate-reader (Infinite 200 Pro, Tecan) at 30 °C. Pepper and sfGFP expression was followed for 3 h measuring fluorescence at 575/620 nm and 485/528 nm respectively. To compare the lysates, end-point values were used for sfGFP and the maximum values for the Pepper aptamer.

nFCM measurement of native *E. coli* vesicles

Native vesicle contents of *E. coli* cell free extracts were measured by nano-flow cytometry (nFCM) using a NanoAnalyzer (NanoFCM Co., Ltd, Nottingham, UK). The device was calibrated with $2.10 \times 10^8$ particles/mL or $2.16 \times 10^8$ particles/mL 250 nm QC beads (NanoFCM Co.) and monodisperse silica beads of four different diameters (68, 91, 113 and 155 nm, NanoFCM Co.) as size reference standards. Samples were diluted in freshly filtered TE buffer which was also measured separately for background subtraction. The measurements were performed for 1 min at a sampling pressure of 1.0 kPa. The NanoFCM software (NF Profession V2.3) was used to calculate particle concentration, mean size and size distribution.

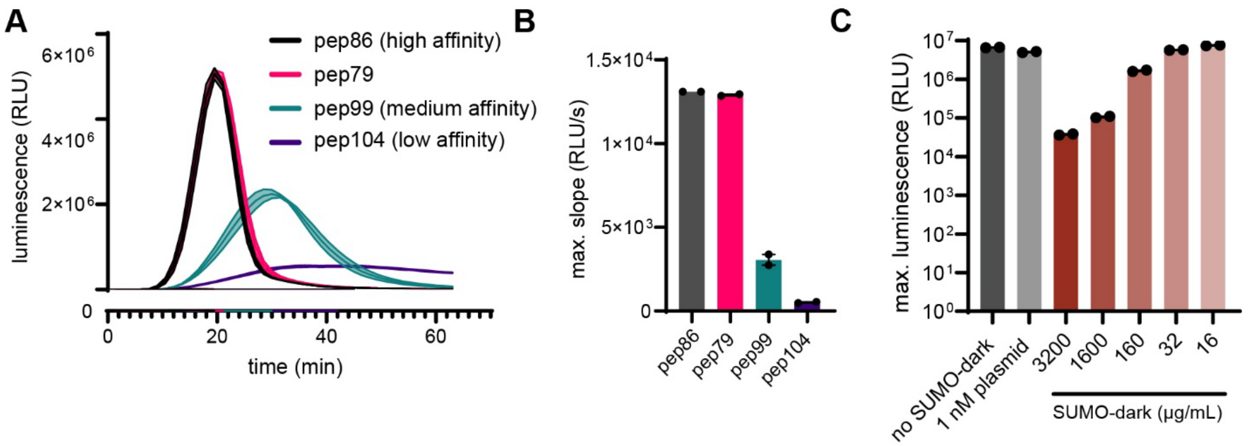


**Fig. S1.**

**Adaption of a split nanoluc system for the use in CFPS.** A) Bulk expression kinetic data of proOmpA reporters in S50 autolysate (11S and furimazine present in bulk solution; no vesicles). proOmpA constructs with high/medium/low affinity complementation tags (pep86/pep99/pep104) with differing binding affinities to 11S were expressed from plasmids. B) Resulting maximum slope data. C) Titration of purified SUMO-dark protein in CFPS with proOmpA-pep86 expressed in bulk.

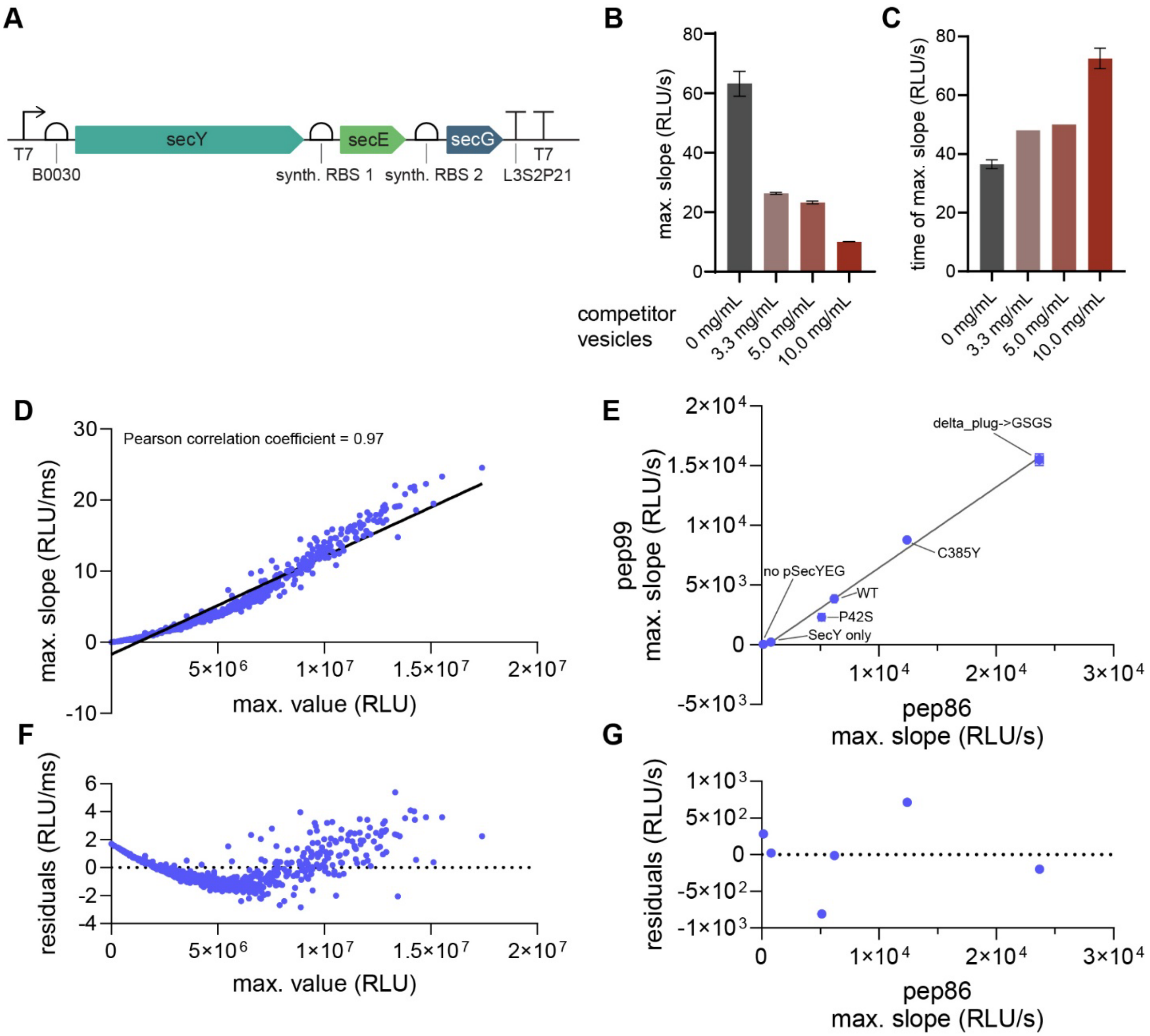


**Fig. S2.**

**Development of the split nanoluc translocation assay for cell free expressed SecYEG.** A) Genetic map of the *secYEG* operon of the pSecYEG construct for cell free expression of functional SecYEG. B) Titration of competitor vesicles (blank membrane; no 11S) in translocation assays using PUREfrex 1.0 and C) their influence on the assay life time. D) Linear regression of maximum luminescence against maximum slope values derived from translocation assay data. F) The respective residuals. Data points were taken from all three translocation replicates of the pSecYEG library. E) Translocation assay data comparing the luminescence output for proOmpA fused to the high (pep86) and medium affinity tag (pep99). Different SecY mutants of pSecYEG were used to compare the two tags. G) Residuals of the linear regression of E).

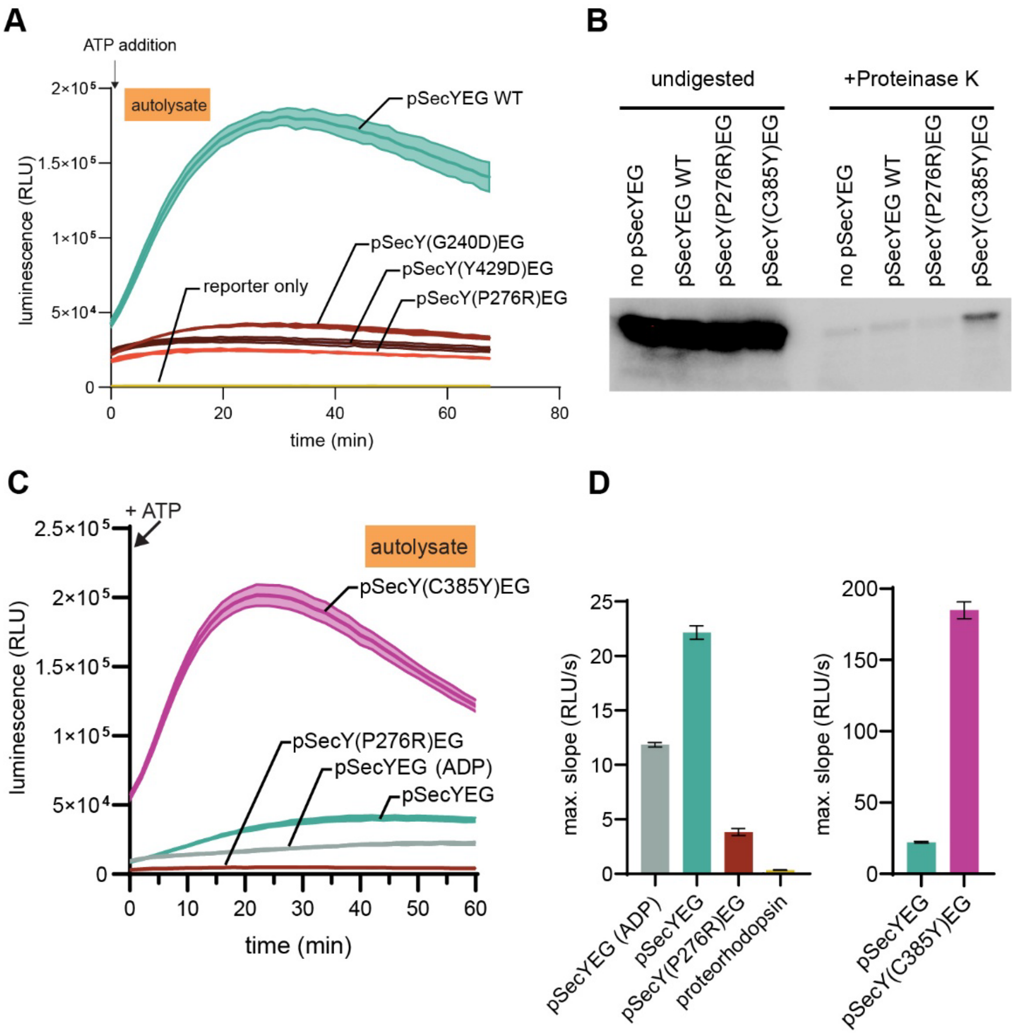


**Fig. S3.**

**Validation of the translocation assay in *E. coli* S50 autolysate.** A) Co-expression of pSecYEG together with the motor ATPase SecA and proOmpA-pep99 in S50 autolysate conveys an ATP dependent translocation signal. 20 mM ATP and 1x furimazine were added after 2.5 h of expressing all three plasmids at the same time. Common translocation deficient SecY mutants (G240D (*20*), Y429D (*20, 21*) and P276R (*19*)) do not respond to ATP addition. B) Protease protection assay of different mutants of the pSecYEG plasmid in lysate. C) Translocation assay similar to a but testing a super-active SecY variants (C385Y (*12, 24*)) and the ATP dependence of SecA driven translocation. D) Maximum slope data of C).

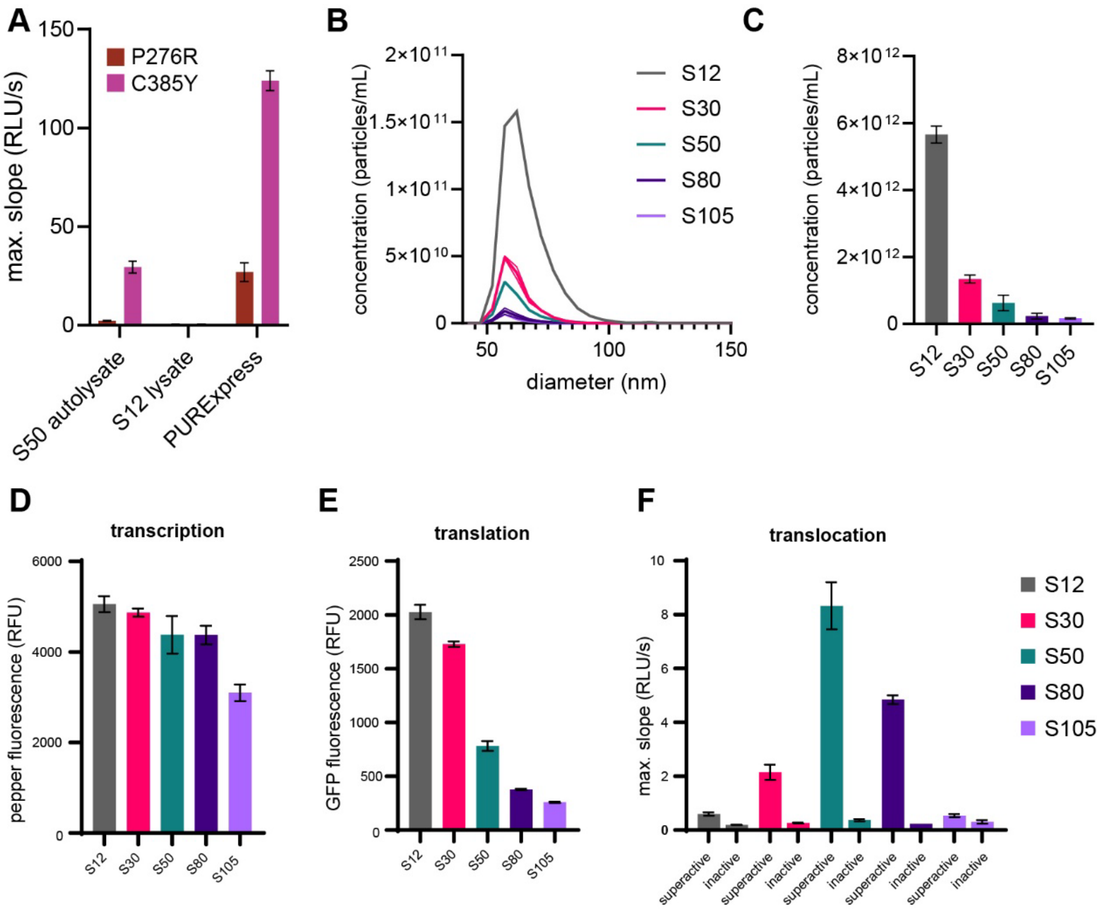


**Fig. S4.**

**SecYEG dependent protein translocation in PURE and cell-free lysates.** A) Translocation assays in three different *E. coli* based CFPS systems using two different variants of pSecYEG with an impaired (P276R (*19*)) and a super-active SecY mutant (C385Y (*12, 24*)). B) Particle size distribution of different five cell free lysates produced by homogenization and cleared at varying centrifugal speeds. C) Total concentration of vesicles in the lysates measured with nFCM. D) Transcription and E) translational activity were assessed using a sfGFP-pepper fusion construct and measuring fluorescence. F) Translocation assay using the C385Y (superactive) and P276R (inactive) SecY mutants.

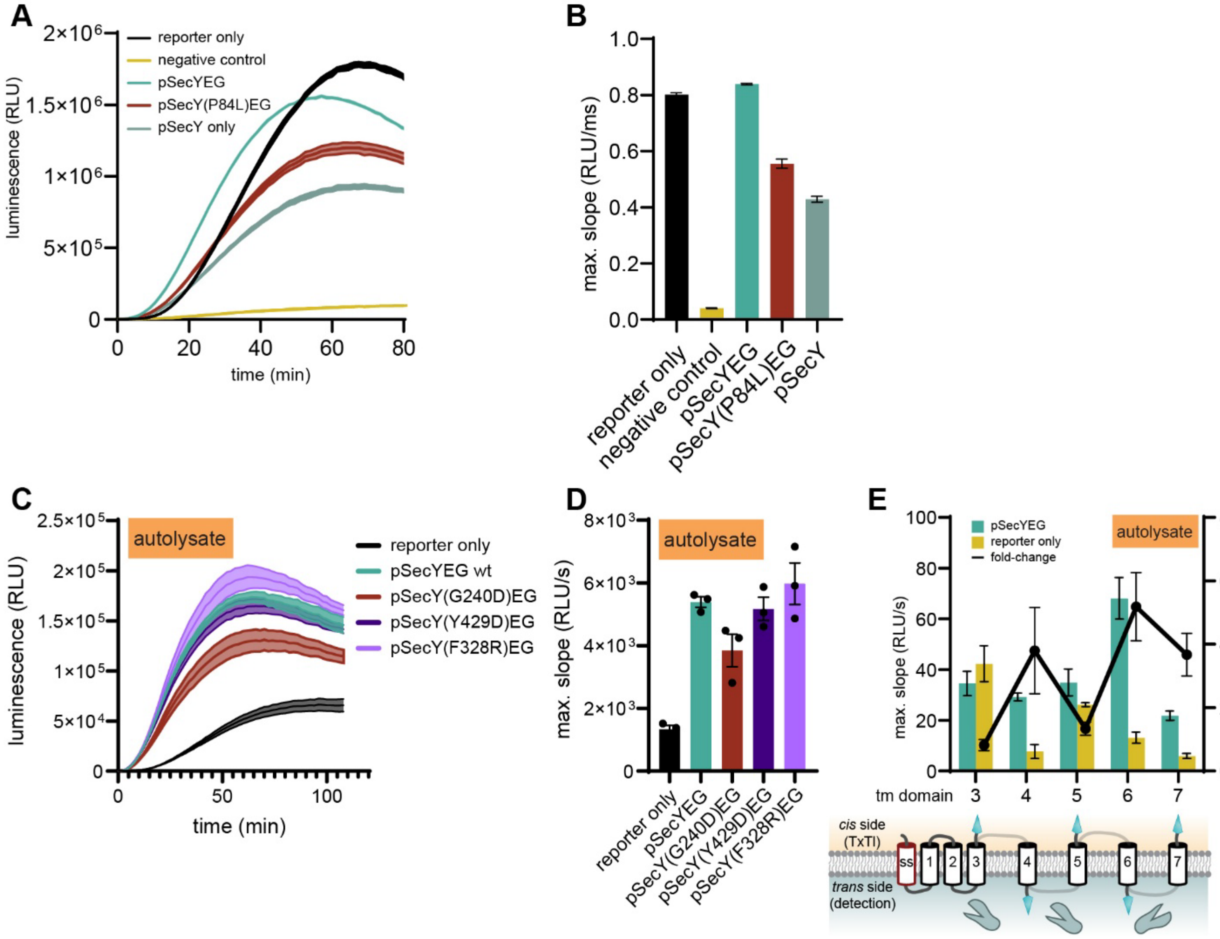


**Fig. S5.**

**Membrane protein insertion assays.** A) Insertion assay in PUREfrex 1.0 using the PR(-1tm)-pep104 reporter. In the "reporter only" control the reporter was expressed alone and without another plasmid compensating for resource competition with pSecYEG. In the negative control, a plasmid encoding native and non-tagged proteorhodopsin was expressed instead of pSecYEG to account for CFPS resource competition. B) Maximum slope values of A). C) Insertion assay in S50 autolysate using the same reporter. The G240D mutant has been reported to be deficient in insertion (*20*) while Y429D has been reported to have to effect on insertion (*20*). D) The respective maximum slope values for different SecY variants of pSecYEG. E) Assessment of directionality of insertion via SecYEG in S50 autolysate.

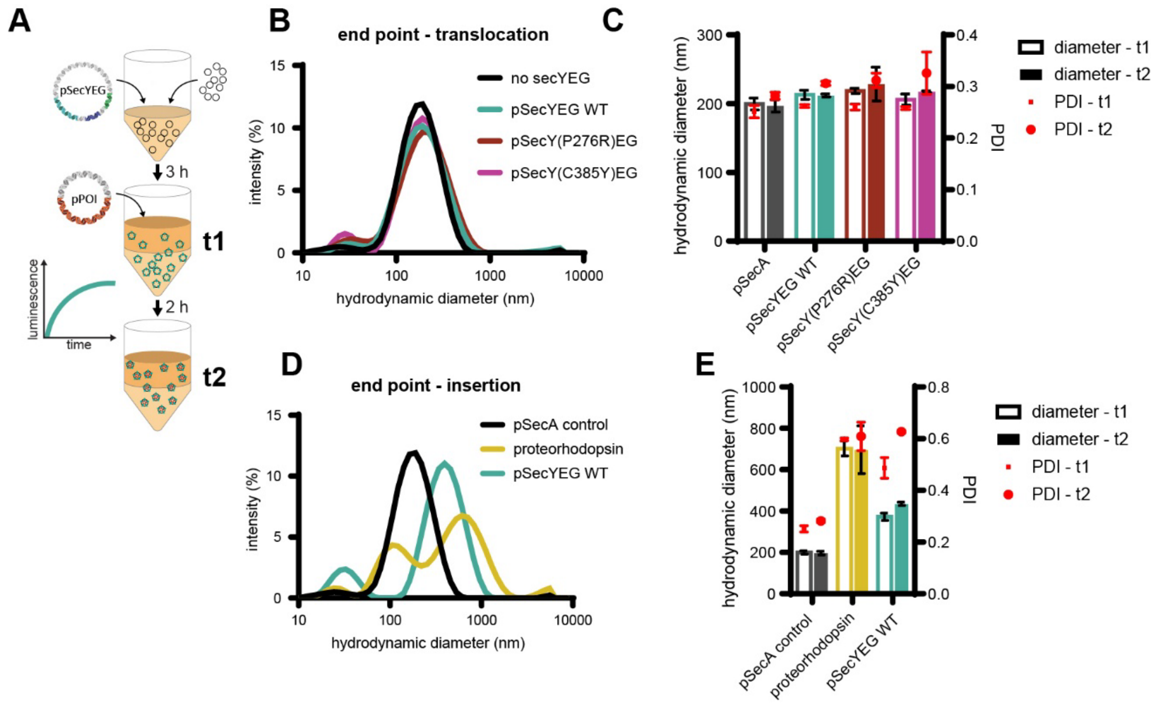


**Fig. S6.**

**Reporter vesicle analysis during translocation and insertion experiments.** A) During standard luminescence translocation (proOmpA-pep99) and insertion (PR(-1tm)-pep104) assays, vesicle samples were analyzed after the first expression phase establishing the SecYEG machinery (t1) and after the reporters had been synthesized (t2). B) DLS analysis at t2 after translocation experiments. C) Mean hydrodynamic diameters and polydispersity indices (PDI) at both time points for translocation. D) DLS analysis at t2 after insertion experiments. E) Mean hydrodynamic diameters and PDI at both time points for insertion.

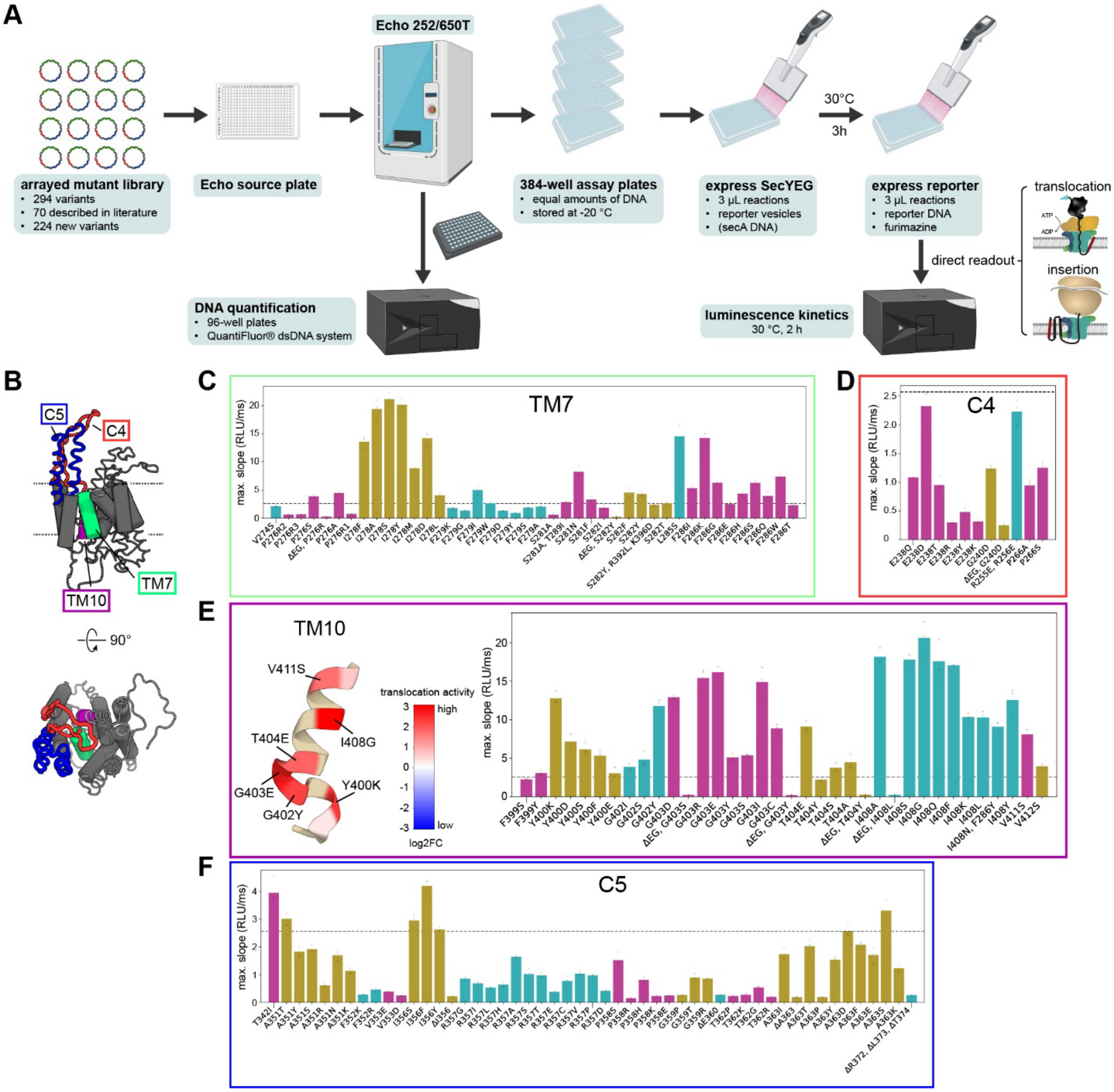


**Fig. S7.**

**Translocation data of the pSecYEG library.** A) Workflow of testing 294 variants of pSecYEG for their translocation and insertion activities (Created in BioRender. Meier, M. (2025) https://BioRender.com/x3y11gh). B) Schematic representation of the SecY structure. Highlighted are the cytoplasmic loops C5 (red) and C5 (blue), and the transmembrane domains TM7 (green) and TM10 (purple) (PDB: 5ABB). The individual translocation signals for each tested mutation within the different domains are shown in C) TM7, D) C4, E) TM10 and F) C5. The dashed horizontal lines indicate the WT activity. For TM10 the mutants with the highest translocation activity for each residue are color coded and plotted on the protein structure.

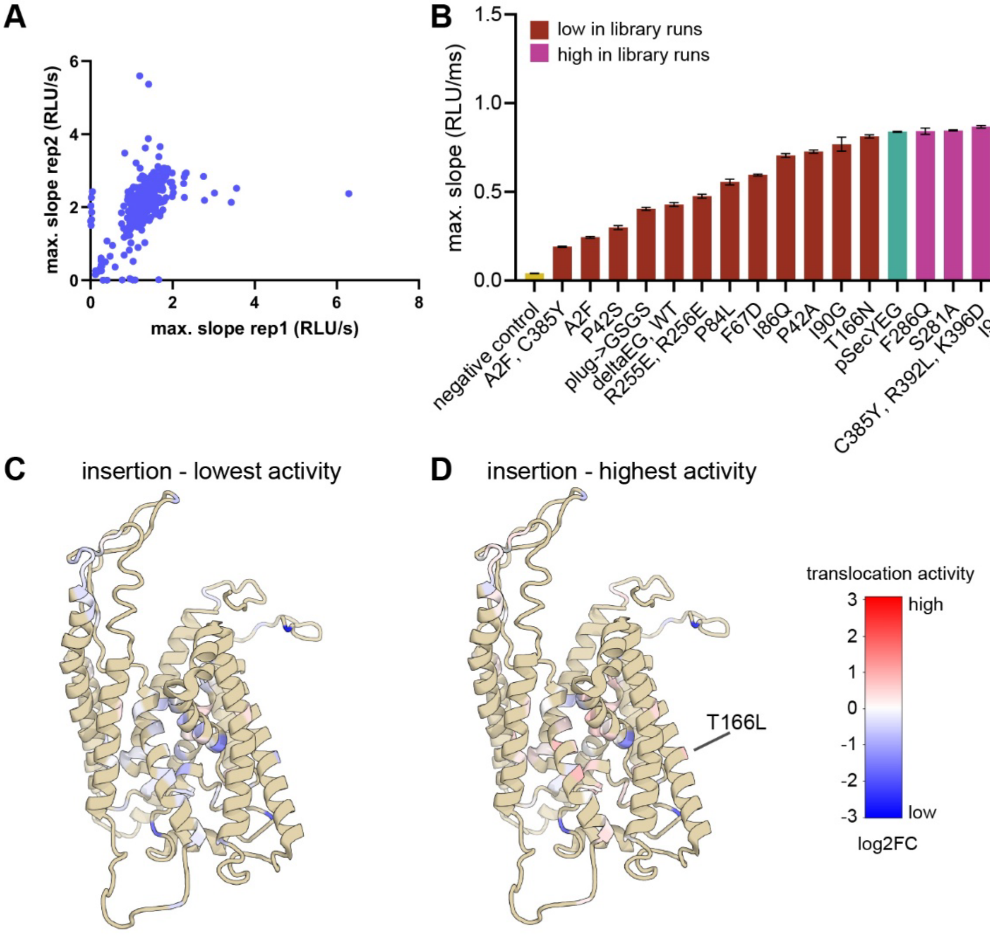


**Fig. S8.**

**Insertion data of the pSecYEG library.** A) Maximum slope values for both replicates of each variant plotted against each other. B) A selection of library plasmids were re-transformed, purified and used in an insertion assay to verify the library results. C) Point mutants with the lowest and D) highest insertion activity obtained for each probed residue are plotted on the SecY structure according to their log2-fold-change over the WT activity (PDB: 5ABB).

**Table S1.**

List of signal peptides used to translocate a nanobody through SecYEG.

| name | organism | UniProt | reference | sequence |
|---|---|---|---|---|
| DsbA | *E. coli* | P0AEG4 | | MKKIWLALAGLVLAFSASA |
| PhoA | *E. coli* | P00634 | | MKQSTIALALLPLLFTPVTKA |
| PpiA | *E. coli* | P0AFL3 | | MFKSTLAAMAAVFALSALSPAAMA |
| TrbC | *E. coli* | P18473 | Kavousipour, 2021 (*59*) | MKLSMKSLAALLMMLNGAVMA |
| YncJ | *E. coli* | P64459 | | MFTKALSVVLLTCALFSGQLMA |
| OmpA | *E. coli* | P0A910 | | MKKTAIAIAVALAGFATVAQAA |
| OmpA(A9N) | *E. coli*, deprecated mutant | | Gennity, 1990 (*49*) | MKKTAIAINVALAGFATVAQAA |
| OmpA(extCore) | synthetic | | Gennity, 1990(*49*) | MKKTAIVAIAIAIATIAVALAGFATVAQAA |
| PorB | *C. glutamicum* | H7C685 | Jeon, 2024 (*47*) | MKLSHRIAAMAATAGITVAAFAAPASA |
| 18F11_Jeon24 | synthetic | | Jeon, 2024 (*47*) | MAGYIRSAFALALAFTAFVFNPPLASA |
| ClustAlign | synthetic | | Kavousipour, 2021 (*59*) | MKKSKALLALALALAASSFALASAFAAFA |

**Data S1. (separate file)**

Extensive translocation and insertion results of the pSecYEG library.

**Data S2. (separate file)**

List of the plasmids used in this study.
